# Acute running improves mood via changes in interoceptive sensibility

**DOI:** 10.64898/2026.06.08.730632

**Authors:** Hideaki Fujihara, Ryuta Kuwamizu

## Abstract

**Purpose:** Interoception is the sensing and interpretation of internal bodily signals and may be a key process linking exercise-induced bodily changes to mood. This study examined whether acute moderate-intensity treadmill running changes interoception and mood states and whether running-related changes in interoception statistically account for mood improvement.

**Methods:** Twenty-seven healthy young adults completed a within-subject crossover experiment consisting of 15 min of moderate-intensity treadmill running at 50% estimated V̇O_₂peak_ and seated rest on separate days. Before and after each condition, cardiac interoceptive accuracy was assessed using heartbeat counting and heartbeat tapping tasks, present-moment interoceptive sensibility using a state-adapted Body Perception Questionnaire (BPQ), and mood using the Profile of Mood States. Running-related changes were calculated as ΔRUN − ΔCTL.

**Results:** Compared with seated rest, running reduced total mood disturbance (TMD) and tension–anxiety (TA) and increased vigor–activity (VA). Running did not significantly change task-based indices of cardiac interoceptive accuracy but increased BPQ scores. Greater running-related increases in BPQ scores were associated with greater reductions in TMD (*r* = −.61, *p* < .001), TA (*r* = −.45, *p* = .018), and depression–dejection (*r* = −.49, *p* = .009). Greater increases in heartbeat tapping task *d′* were associated with greater increases in VA (*r* = .56, *p* = .003). Mediation analysis indicated that BPQ changes were statistically consistent with an indirect pathway linking running to reduced TMD, with weaker exploratory evidence for TA.

**Conclusions:** A single bout of moderate-intensity running improved mood and increased present-moment interoceptive sensibility, but not task-based cardiac interoceptive accuracy. Enhanced subjective awareness of bodily sensations may be one proximal process related to the mood benefits of acute running.

## Introduction

Acute aerobic exercise has been widely studied as a strategy for improving mood, particularly by reducing anxiety. Previous studies have reported that even a single bout of exercise can significantly reduce state anxiety relative to pre-exercise levels (1). Recent meta-analytic evidence has also shown that moderate- to high-intensity exercise is associated with reduced anxiety in healthy adults (1,2,3). However, not all individuals benefit from acute exercise to the same extent (3), suggesting that the anxiolytic effects of exercise may not be uniform across individuals. Therefore, to establish exercise more firmly as a strategy for promoting mental health and reducing anxiety-related mood disturbances, it is necessary to clarify the proximal mechanisms underlying exercise-induced reductions in anxiety, as well as the factors that contribute to individual differences in these effects.

A growing body of physiological evidence has identified several neurological mechanisms through which exercise may improve mood states, including the regulation of monoaminergic systems and the hypothalamic–pituitary–adrenal axis, the release of endogenous opioids and endocannabinoids, and the expression of neurotrophic factors such as brain-derived neurotrophic factor (BDNF) (4,5,6,7). Psychological mechanisms, such as enhanced self-efficacy and attentional distraction, have also been proposed (8,9). However, these mechanisms explain only certain aspects of the complex responses induced by exercise, and it remains difficult to account for, within a single pathway, the proximal process by which exercise-induced changes in bodily states are perceived, evaluated, and ultimately linked to subjective changes in mood states (10). These considerations raise the possibility that individual differences in the mood-enhancing effects of exercise may depend not only on the magnitude of physiological changes induced by exercise, but also on how these bodily changes are perceived, interpreted, and integrated into subjective emotional experience.

Accordingly, the present study focused on interoception. Interoception is defined as the process by which the nervous system senses, interprets, and integrates internal signals from bodily systems, such as the cardiovascular, respiratory, and gastrointestinal systems, to form representations of bodily states (11,12). From a neuroanatomical perspective, interoceptive processing is supported by a network centered on the insular cortex (insula) and anterior cingulate cortex (ACC), both of which are deeply involved in emotion generation and subjective emotional experience (11,13,14). In particular, the integration of bodily signals in the insula has been associated with subjective anxiety (14,15), suggesting that interoception may constitute a fundamental mechanism linking bodily states to emotional experience.

Acute aerobic exercise produces salient changes in internal bodily states, including increases in heart rate (HR) and respiration. Such exercise-induced bodily signals are thought to engage interoceptive pathways involving afferent feedback from the body and brain regions such as the insula and ACC, thereby influencing cognitive, motivational, and emotional processes (16). Such changes may make bodily signals more noticeable and alter how individuals perceive and attend to their bodily states, thereby leading to changes in interoceptive processing (17). Importantly, anxiety has been linked to altered interoceptive processing, particularly heightened attention to bodily sensations, negative evaluation of these sensations, and difficulties describing bodily signals and emotions (18,19). Intentional activities such as exercise may provide a context in which bodily signals are generated in a relatively clear and predictable manner. By increasing the salience and interpretability of bodily signals, exercise may alter interoception and facilitate the integration of bodily changes into subjective emotional experience, thereby contributing to changes in mood states (12,20). Indeed, previous studies have suggested that acute physical activity can improve performance on heartbeat perception tasks (e.g., Ref. 21), possibly by increasing the salience of cardiac signals (22,23). However, it remains unclear whether exercise-induced changes in interoception are associated with changes in mood states, and whether interoception can account for individual differences in mood improvement after acute exercise. In particular, few studies have tested the exercise–interoception–mood pathway using a mediation model.

Based on this background, the present study examined the effects of acute moderate-intensity aerobic exercise, specifically treadmill running, on mood states and interoception in healthy university students. Moderate-intensity exercise is commonly recommended for health promotion (24), and the present study operationalized moderate-intensity running as 50% of estimated peak oxygen uptake (V̇O₂_peak_). Treadmill running was selected because recent evidence suggests that the effects of exercise on mood may differ depending on exercise modality. For example, several studies have reported that cycling exercise increases arousal, whereas its effects on pleasant mood are relatively limited (25,26,27). In contrast, running has been shown to increase both arousal and pleasant mood (28,29). Although the mechanisms underlying these modality-specific effects remain unclear, one possible explanation is that running-specific mechanical stimulation, such as head acceleration, may contribute to changes in mood and cognition (29,30). Therefore, the present study hypothesized that changes in interoception induced by moderate-intensity running would contribute to reductions in negative mood states and broader improvements in mood. We tested this hypothesis using a quasi-randomized crossover design and examined whether running-related changes in interoception were associated with, and statistically accounted for, changes in mood states.

## Methods

### Participants

Thirty healthy young adults participated in this study. All participants were native Japanese speakers, were right-handed, and had normal or corrected-to-normal vision and normal color vision. No participants reported taking medication affecting the central nervous system or endocrine system. Participants were excluded if they had severe depressive symptoms, defined as a Beck Depression Inventory-II (BDI-II) score of 29 or higher (n = 1), or if they did not follow the task instructions correctly (n = 2). Consequently, 27 participants (14 female) were included in the final analysis. Participant characteristics are shown in Table 1.

**Table 1.**
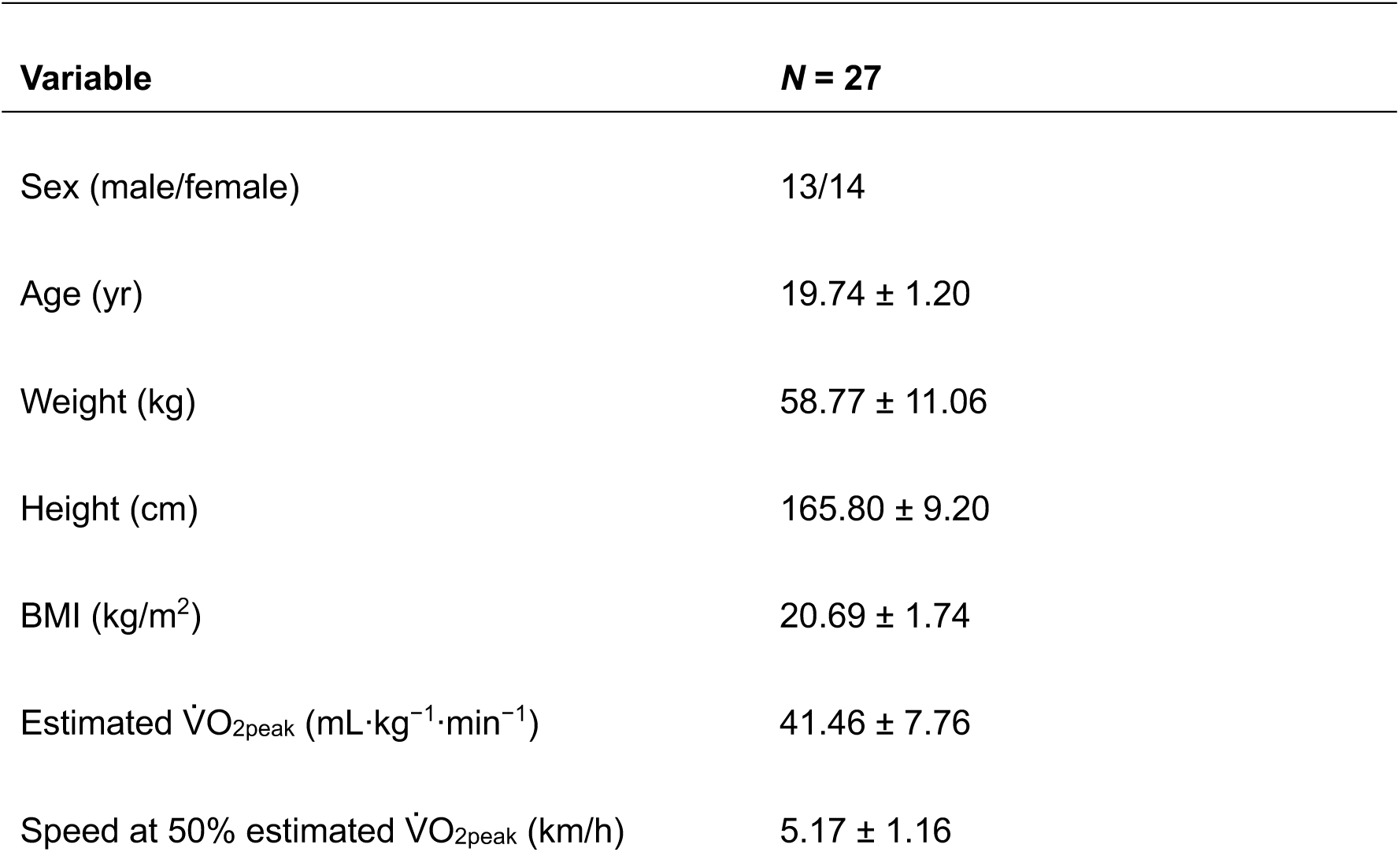
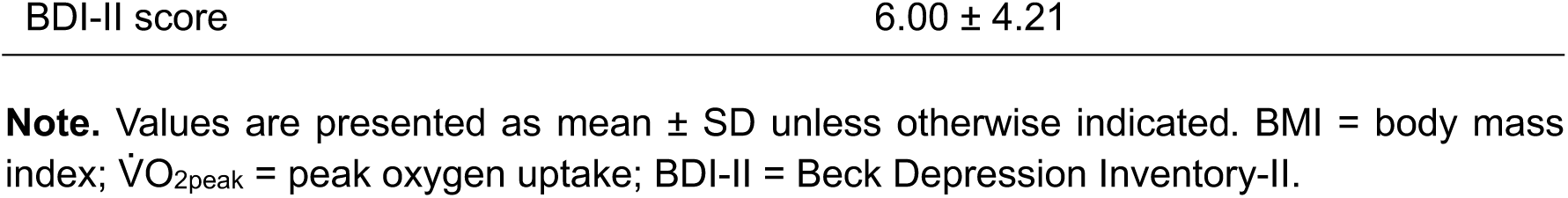
Participant characteristics.

The final sample size was considered acceptable for the present within-subject crossover design. Based on Green’s guideline (31), at least 31 observations are required for multiple regression analysis with three predictors when assuming a large effect size (f² > 0.35), as used in the causal mediation analyses. In the present study, 27 participants completed both the running and control conditions, yielding 54 condition-level observations for the mediation analyses.

This study was approved by the Institutional Ethics Committee of the Graduate School of Social and Industrial Science and Technology, Tokushima University (approval number 367). All procedures were conducted in accordance with the Declaration of Helsinki. Complete information about the study was provided to participants, and written informed consent was obtained from all participants before participation.

### Experimental design and procedure

The study consisted of a baseline assessment day and two main experimental sessions. On Day 1, participants completed the single-stage submaximal treadmill walking test (SSTWT) to estimate maximal oxygen uptake and determine the individualized treadmill workload corresponding to 50% of estimated peak oxygen uptake (V̇O_2peak_) for the RUN condition. On the same day, participants also completed the Body Perception Questionnaire–Body Awareness Very Short Form, Japanese version (BPQ-BAVSF-J), BDI-II, and anthropometric measurements of height and weight.

The main experiment consisted of two conditions: a moderate-intensity running condition (RUN) and a seated resting control condition (CTL). All participants completed both conditions on separate days after the baseline assessment session. The order of the RUN and CTL conditions was quasi-randomized across participants to reduce potential order effects. The two main experimental sessions were separated by at least 48 h.

To control for external influences, participants were instructed to refrain from exercise and alcohol consumption for at least 24 h before each main experimental session and to avoid caffeine on the day of each experimental session. Interoceptive measures and mood states were assessed immediately before and after each condition using the same task order and procedure across conditions. The overall experimental procedure is shown in Figure 1.

**Figure 1.**
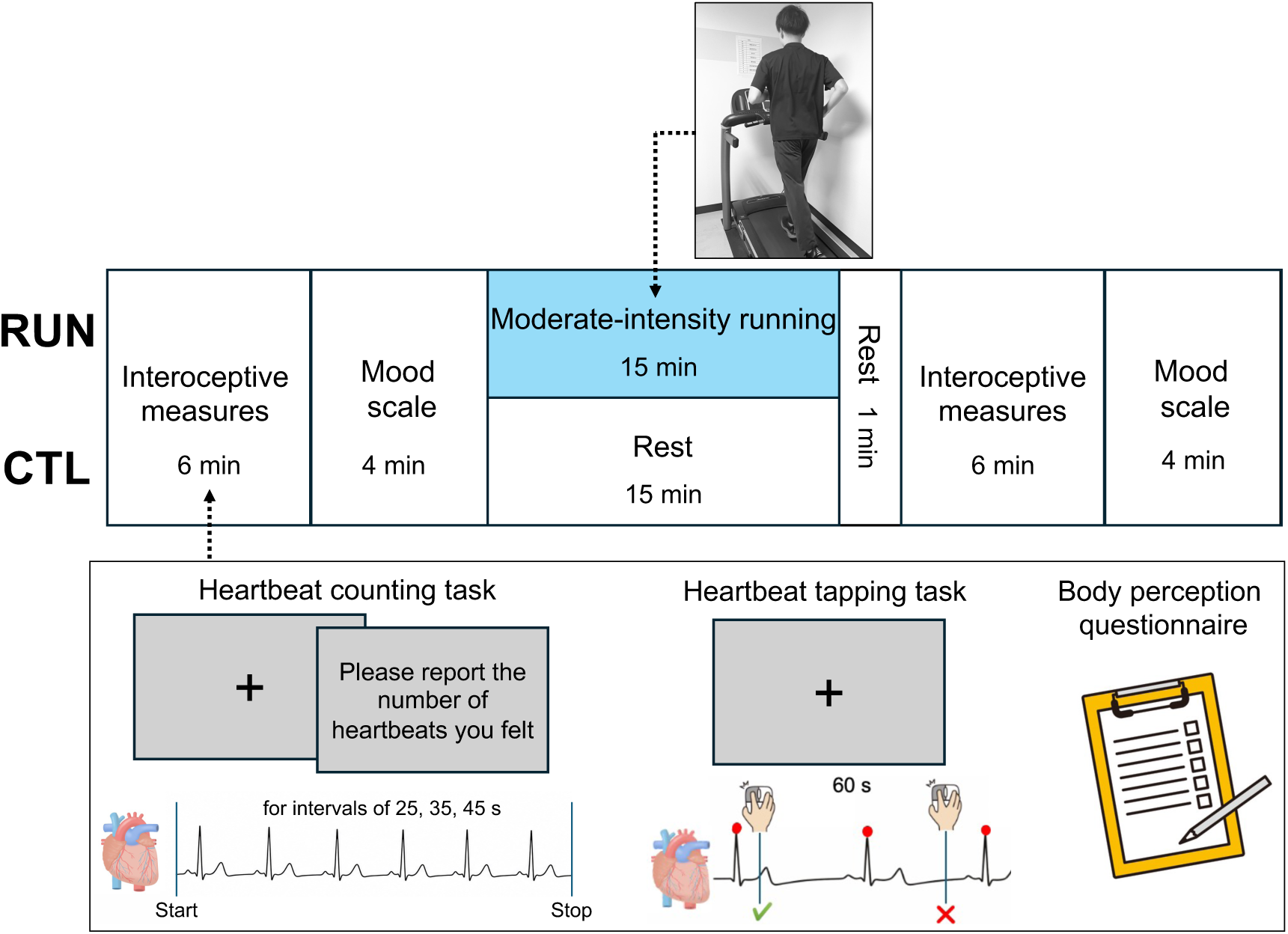
Experimental design for testing acute running effects on interoception and mood. Interoceptive measures and mood were assessed immediately before and after each condition. In the RUN condition, participants completed 15 min of moderate-intensity treadmill running, whereas in the CTL condition, they remained seated quietly for 15 min. Interoceptive measures included the Heartbeat counting task (HCT), Heartbeat tapping task (HTT), and Body perception questionnaire (BPQ). In the HCT, participants silently counted perceived heartbeats during intervals of 25, 35, and 45 s and then reported the number. In the HTT, participants clicked a mouse button whenever they felt a heartbeat during a 60-s trial. Both tasks were preceded by a 10-s practice trial. The BPQ assessed self-reported present-moment interoceptive sensibility. Note: RUN = moderate-intensity running condition; CTL = seated resting control condition.

### Single-stage submaximal treadmill walking test

Participants performed an SSTWT to estimate V̇O_2peak_ and to determine an individualized moderate-intensity exercise workload corresponding to 50% intensity, in accordance with the guidelines of the American College of Sports Medicine (32). The SSTWT, originally developed by Ebbeling and colleagues, is a widely used method for estimating V̇O_2peak_ (33), as previous studies have demonstrated a strong association between predicted and directly measured V̇O_2peak_ values (*r* = 0.96, with a multiple correlation 0.86)(34). The test consisted of two consecutive 4-minute stages. During the initial stage, each participant walked on a treadmill at a self-selected speed ranging from 2.0 to 4.5 mph with no incline, serving as a warm-up period. Then, treadmill speed was gradually adjusted until the participant’s heart rate reached approximately 60–70% of their age-predicted maximal heart rate (HR_max_), calculated using the equation 207 − (0.67 × age). Throughout the second 4-min stage, participants were required to maintain the speed they had established during the first stage while the treadmill was raised to a 5% grade. Steady-state heart rate (SSHR) was calculated as the average heart rate during the last 2 min of the second stage and, together with age, sex, and final treadmill speed, was used to estimate V̇O₂_peak_ using the following equation: V̇O_2peak_ (mL/kg/min) = 15.1 + 21.8 × speed (mph) − 0.327 × SSHR (bpm) − 0.263 × speed × age (years) + 0.00504 × SSHR × age + 5.98 × sex (32), where speed indicates the final treadmill speed in mph and sex is coded as 0 for female and 1 for male. Then, based on the ACSM metabolic equation for treadmill running (35), the treadmill running speed corresponding to 50% of the estimated V̇O_2peak_ at 0% incline was individually determined for each participant (Table 1). This intensity corresponds to the moderate-intensity range according to the ACSM guidelines for exercise prescription (36).

### Exercise and control conditions

In the RUN condition, participants performed 15 min of treadmill running at an individualized workload corresponding to 50% of each participant’s estimated V̇O_2peak_ at 0% incline. This intensity was selected because 50% V̇O_2peak_ is classified as moderate intensity according to the American College of Sports Medicine (36). The exercise duration was set at 15 min because previous studies have suggested that at least 10–20 min of running can improve mood (28,37,38).

During the RUN condition, HR and RPE were monitored every minute to verify exercise intensity. In the CTL condition, participants remained seated quietly on a chair for 15 min. The duration of the CTL condition was matched to that of the RUN condition.

Interoceptive measures and mood states were assessed immediately before and after both the RUN and CTL conditions using the same procedure and task order.

### Interoceptive accuracy tasks

#### Heartbeat counting task (HCT)

The HCT used in the present study was based on the Mental Tracking Method originally proposed by Schandry (39). Participants completed three trials with intervals of 25, 35, and 45 s, presented in a randomized order across participants. A 20-s rest period was provided between trials. The beginning and end of each trial were signaled by the appearance and disappearance of a fixation point presented at the center of the screen.

Participants were instructed to remain seated quietly and to count their own heartbeats as accurately as possible during each interval, even if they were uncertain about their responses. After each trial, they reported the number of counted heartbeats by entering the value on an on-screen numeric keypad using a mouse, with no time limit for responding. Before the experimental trials, one 10-s practice trial was administered to ensure that participants understood the task instructions and became familiar with the procedure.

#### Heartbeat tapping task (HTT)

Although the HCT is widely used to assess interoceptive accuracy, its performance can be influenced by non-interoceptive factors, such as prior knowledge or beliefs about one’s own heart rate and estimation strategies, rather than solely by the perception of actual heartbeat sensations (40,41,42). Therefore, in addition to the HCT, the present study included an HTT, which requires participants to tap in synchrony with their perceived heartbeats and allows the temporal correspondence between cardiac events and behavioral responses to be assessed. The design of this task was based on similar procedures reported in previous studies (17,43,44,45,46).

A fixation point was continuously presented at the center of the screen throughout each trial. Participants were instructed to respond by clicking the left mouse button in synchrony with their own heartbeat as accurately as possible while the fixation point was displayed. Even when they were uncertain, they were asked to respond based on their best estimate. This instruction is consistent with standard procedures commonly used in heartbeat perception tasks (17,47). Before the experimental trial, one 10-s practice trial was administered to facilitate understanding of the task procedure. After the practice trial, participants performed one 60-s experimental trial.

### Interoceptive sensibility

Interoceptive sensibility was assessed using the Japanese version of the Body Perception Questionnaire (BPQ) –Body Awareness Very Short Form. The BPQ was originally developed in English as a measure of bodily awareness (48,49). In the present study, we used the validated Japanese version of the BPQ-BAVSF (50), a concise 12-item scale designed to assess individual differences in sensitivity to internal bodily sensations and physiological functions. The items cover bodily experiences related to respiration, cardiovascular activity, muscle tension, gastrointestinal processes, and other internal bodily cues. Responses are rated on a 5-point Likert scale ranging from “Never” to “Always.” Total scores range from 12 to 60, with higher scores indicating greater sensitivity to bodily sensations. For clarity and brevity, scores derived from the BPQ-BAVSF-J are referred to as “BPQ scores” throughout this study.

The standard BPQ-BAVSF-J is administered in a trait-based format, asking respondents to rate their awareness of bodily sensations during most situations. However, because the present study aimed to examine state-related changes in interoceptive sensibility before and after the experimental sessions, the wording of the instructions was modified with permission from the first author of the Japanese version (50). In the modified version used in this study, participants were asked to rate each item according to their present-moment bodily experience at the time of assessment. For example, an item such as “My mouth being dry” was rated with reference to the participant’s current bodily state. This adaptation was intended to allow the BPQ-BAVSF-J to capture short-term fluctuations in interoceptive sensibility associated with the running intervention.

To examine whether this state-based adaptation retained continuity with the original trait-based construct, participants completed the original trait version of the BPQ-BAVSF-J on Day 1. Because the RUN and CTL sessions took place on either Day 2 or Day 3 depending on the participant, we examined correlations between the modified BPQ scores obtained at the pre-assessment prior to each session on Day 2 and Day 3 and the original baseline BPQ scores obtained on Day 1. The modified BPQ scores were significantly positively correlated with scores on the original version on both Day 2 and Day 3 (Day 2: *r* = .50, *p* = .009; Day 3: *r* = .56, *p* = .002). These correlations provide preliminary support that the modified version retained continuity with the construct measured by the original trait-based BPQ-BAVSF-J. Given the state-adapted nature of the modified instructions, BPQ scores in the present study are described as self-reported present-moment interoceptive sensibility.

### Mood states

Mood states were assessed using the Japanese version of the Profile of Mood States 2nd Edition–Adult Short Form (POMS 2-A Short) (51,52). The POMS 2-A Short is a 35-item self-report instrument designed to evaluate transient mood states. It yields scores on seven mood dimensions: anger–hostility (AH), confusion–bewilderment (CB), depression–dejection (DD), fatigue–inertia (FI), tension–anxiety (TA), vigor–activity (VA), and friendliness (F). Participants rated each item on a 5-point Likert scale ranging from 0 (“not at all”) to 4 (“extremely”).

Higher scores on the VA and F subscales reflect more positive mood states, whereas higher scores on the remaining five subscales indicate greater severity of negative mood symptoms. In addition to the subscale scores, an overall index of mood disturbance, the TMD score, was computed. The total mood disturbance (TMD) score was calculated as AH + CB + DD + FI + TA − VA, with higher values representing greater TMD.

### Electrocardiographic measurement

Electrocardiographic (ECG) signals were recorded using biosignalsplux acquisition devices (PLUX Wireless Biosignals, Lisbon, Portugal) at a sampling rate of 1000 Hz to continuously assess the electrical timing of participants’ heartbeats throughout the interoceptive accuracy tasks. The ECG sensor’s three-electrode montage, including ground, was attached to the left side of the chest, with two recording electrodes approximately positioned over the V4–V5 region and the ground electrode placed inferior to them to ensure stable signal acquisition.

ECG and behavioral response data were processed offline using MATLAB code developed in-house. R-peaks were detected automatically from the ECG signal, and the detected peaks were subsequently visually inspected to verify the absence of obvious artifacts, outliers, or abnormal detections.

During the SSTWT and the RUN session, instantaneous HR was monitored and recorded using a myBeat device with ECG electrodes (UNION TOOL, Tokyo, Japan) placed over the V1–V2 region. This recording was used to monitor participants’ heart rate responses during exercise and to ensure that the exercise intensity was appropriately maintained throughout the treadmill protocol.

### Perceptual measures

The HCT score was calculated across the three trials using the following formula: HCT score = 1/3 Σ (1 − |actual number of heartbeats − self-reported number of heartbeats| / actual number of heartbeats). Higher scores were interpreted as indicating greater interoceptive accuracy.

For the HTT, perceptual performance was evaluated using three indices adapted from the general framework proposed by Fittipaldi et al. (44): mean distance (md), modified Schandry index (mSI), and *d′*. The md index quantifies the discrepancy between cardiac frequency and tapping frequency across overlapping time windows, with lower values indicating closer correspondence between heartbeat rhythm and tapping rhythm. The mSI quantifies the correspondence between the total number of detected R-peaks and the total number of taps, with higher values indicating greater response-count agreement. The *d′* index, based on Signal Detection Theory, quantifies participants’ sensitivity in discriminating heartbeat-related signals from non-signal responses, with higher values indicating greater sensitivity. Detailed calculation procedures for these indices are provided in Supplemental Methods (see Supplemental Methods, Supplemental Digital Content 1, which provides detailed calculation procedures for the HTT-derived md, mSI, and *d′* indices).

### Statistical analyses

Statistical analyses were performed in R version 4.3.2. Interoceptive accuracy, interoceptive sensibility, and mood scores were analyzed using repeated-measures two-way ANOVAs with condition (RUN, CTL) and time (Pre, Post) as within-subject factors. These analyses were conducted using the afex package (version 1.4.1), with Type III sums of squares and partial eta squared (*ηp*²) reported as the effect size. When significant condition × time interactions were observed, Bonferroni-corrected post hoc comparisons were conducted using the emmeans package (version 1.11.2). In addition, for complementary purposes, change scores for each condition were calculated as Post − Pre, and paired *t*-tests were used to compare ΔRUN and ΔCTL. To examine the association between running-related changes in interoception and mood, contrast scores representing running-related effects were computed as ΔRUN − ΔCTL, and Pearson’s correlation analyses were conducted between contrast scores for interoceptive measures and those for POMS outcomes. All statistical tests were two-sided, and statistical significance was set a priori at *p* < .05.

Mediation analyses were performed using the mediation package (version 4.5.0) (53) to examine whether running-related changes in interoceptive accuracy or present-moment interoceptive sensibility were statistically consistent with an indirect pathway linking running to mood improvement. Because the present study used a within-subject crossover design, mediation analyses were conducted using fixed-effects linear models with participant ID included as a fixed effect. For each participant and condition, condition-specific change scores were calculated as Post − Pre. The exposure variable was condition, coded as 1 for RUN and 0 for CTL. In each mediation model, the mediator was defined as the condition-specific change in an interoceptive measure, and the outcome was defined as the condition-specific change in the corresponding mood score. The mediator model included condition and participant fixed effects, and the outcome model included condition, the mediator, and participant fixed effects. The total effect, average direct effect (ADE), average causal mediation effect (ACME), and proportion mediated were estimated using quasi-Bayesian confidence intervals based on 5,000 simulations. Candidate mediation models were selected based on two criteria: first, both the interoceptive measure and the mood outcome showed significant condition × time interactions, indicating running-related changes; second, running-related changes in the interoceptive measure were significantly correlated with running-related changes in the mood outcome. TMD was specified as the primary mood outcome, and POMS subscales meeting these criteria were treated as exploratory outcomes. Because changes in the mediator and outcome were assessed over the same pre–post interval, these mediation analyses were interpreted as preliminary and as testing whether the data were statistically consistent with an indirect pathway, rather than as providing definitive evidence of causal mediation.

## Results

### Verification of moderate-intensity running

We first verified that the RUN condition induced moderate-intensity physiological responses. HR and rating of perceived exertion (RPE) were monitored every minute during the running session (54). During the last minute of running, average HR and RPE were 141.28 ± 13.34 bpm and 12.63 ± 1.80, respectively. These values correspond to the moderate-intensity range according to ACSM-based exercise intensity criteria (36). We further examined whether the RUN condition led to increased cardiac activity at the time of the subsequent interoceptive accuracy task. A two-way repeated-measures ANOVA on the number of R-peaks recorded during the 60-s HTT revealed a significant condition × time interaction (*F* (1, 26) = 68.33, *p* < .001, *ηp*² = .724). Post hoc comparisons showed that R-peak counts significantly increased between pre and post in the RUN condition (75.22 ± 9.99 to 89.70 ± 11.58; *t* (26) = −7.83, *p* < .001, *d* = −1.51). At post, R-peak counts were significantly higher in the RUN condition than in the CTL condition (89.70 ± 11.58 vs. 71.89 ± 11.60; *t* (26) = 7.88, *p* < .001, *d* = 1.52). These findings indicate that moderate-intensity running led to increased cardiac activity at the time of the post-exercise interoceptive accuracy task.

### Interoceptive accuracy and sensibility

We next examined whether moderate-intensity running altered objective and subjective aspects of interoception. Interoceptive accuracy was assessed using heartbeat perception tasks. Present-moment interoceptive sensibility was assessed using BPQ scores. Changes in these measures before and after the RUN and CTL conditions are shown in Figure 2.

**Figure 2.**
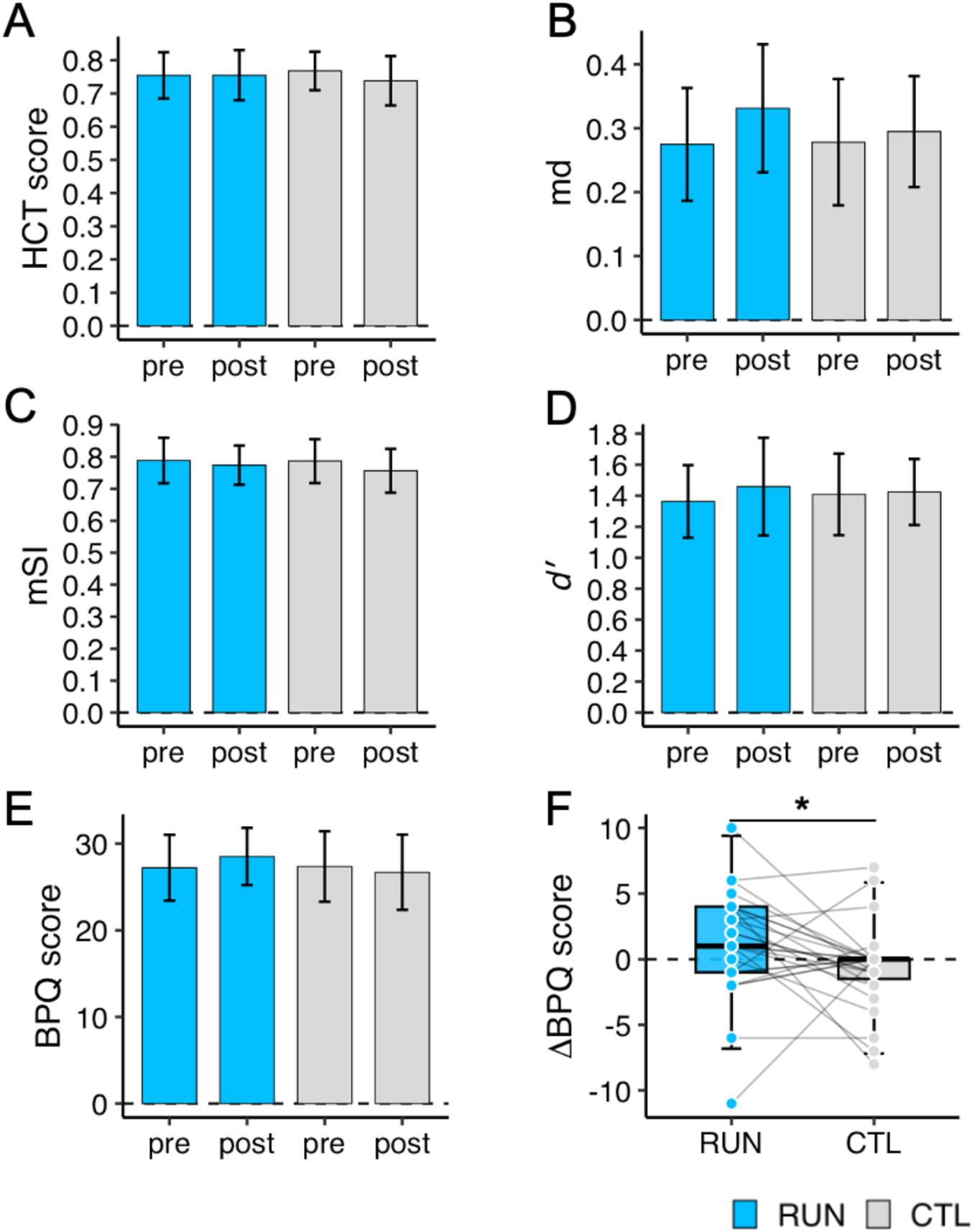
Effects of running on interoceptive accuracy indices and present-moment interoceptive sensibility. (A) HCT scores before and after the RUN and CTL conditions. (B) HTT mean distance (md) before and after the RUN and CTL conditions. Lower md values indicate closer correspondence between heartbeat rhythm and tapping rhythm. (C) HTT modified Schandry index (mSI) before and after the RUN and CTL conditions. (D) HTT *d′* scores before and after the RUN and CTL conditions. (E) BPQ scores before and after the RUN and CTL conditions. BPQ scores were used as an index of self-reported present-moment interoceptive sensibility. (F) Changes in BPQ scores for the RUN and CTL conditions. The increase in BPQ score was significantly greater in the RUN condition than in the CTL condition, suggesting that moderate-intensity running enhanced self-reported present-moment interoceptive sensibility. **Note**: \**p* < 0.05. In panels A–E, bars represent means and error bars indicate 95% confidence intervals. In panel F, box-and-whisker plots are drawn in the Tukey manner, overlaid error bars indicate mean ± 2 SD, dashed horizontal lines indicate zero, and line plots represent individual data. RUN = moderate-intensity running condition; CTL = seated resting control condition; HCT = heartbeat counting task; HTT = heartbeat tapping task; md = mean distance; mSI = modified Schandry index; BPQ = Body Perception Questionnaire.

For the interoceptive accuracy indices, repeated-measures two-way ANOVAs with condition (RUN, CTL) and time (Pre, Post) as within-subject factors revealed no significant condition × time interactions for HCT score, HTT md, HTT mSI, or HTT *d′* (see Supplemental Table 1, Supplemental Digital Content 1, which presents repeated-measures ANOVA results for interoceptive accuracy indices and present-moment interoceptive sensibility; Figure 2A–D). These findings indicate that moderate-intensity running did not significantly alter task-based measures of interoceptive accuracy in the present study.

For BPQ scores, the repeated-measures two-way ANOVA also revealed a significant condition × time interaction (*F* (1, 26) = 4.29, *p* = .048, *ηp*² = .14). However, Bonferroni-corrected post hoc comparisons did not reveal significant differences between pre- and post-session scores within either condition or between conditions at either time point (Figure 2E). In the complementary change-score analysis, the increase in BPQ score was significantly greater in the RUN condition than in the CTL condition (*t* (26) = 2.07, *p* = .048, *d* = 0.40)(Figure 2F). These findings suggest that moderate-intensity running increased self-reported present-moment interoceptive sensibility compared with seated rest.

### Mood states

We next examined whether moderate-intensity running improved mood states assessed using the POMS 2-A Short. Figure 3 A shows changes in TMD scores before and after the RUN and CTL conditions. For TMD score, a repeated-measures two-way ANOVA with condition (RUN, CTL) and time (Pre, Post) as within-subject factors revealed a significant condition × time interaction, *F* (1, 26) = 11.95, *p* = .002, *ηp*² = .32. Bonferroni-corrected post hoc comparisons showed that TMD score significantly decreased between the pre- and post-sessions in the RUN condition (*t* (26) = 3.39, *p* = .002, *d* = 0.65), indicating that moderate-intensity running reduced TMD. Consistent with this interaction, TMD decreased significantly more in the RUN condition than in the CTL condition (*t* (26) = −3.46, *p* = .002, *d* = −0.67)(Figure 3B), indicating a greater reduction in TMD after running than after seated rest.

**Figure 3.**
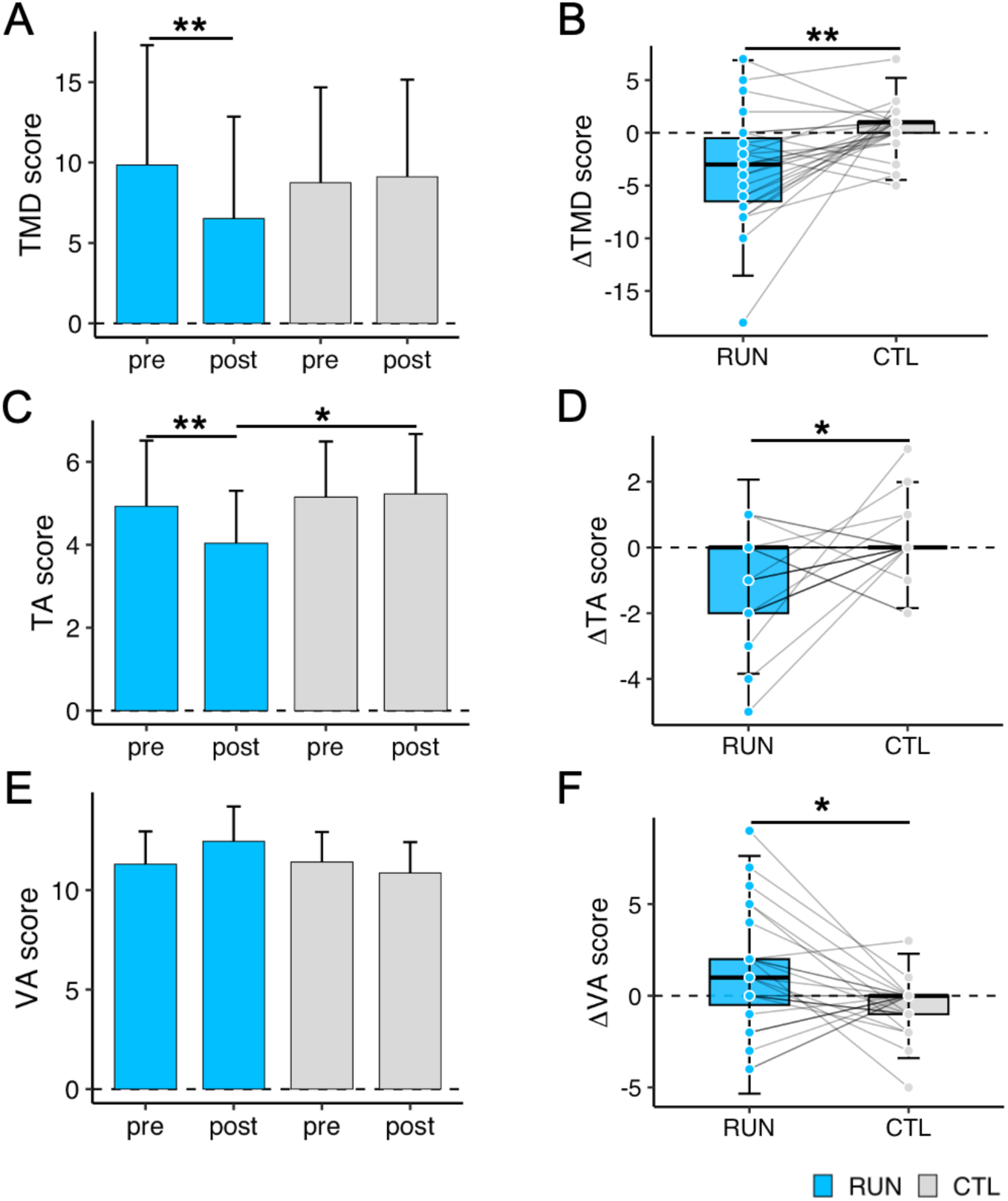
Effects of running on mood states. (A) TMD scores before and after the RUN and CTL conditions. Moderate-intensity running led to significantly reduced TMD scores between pre- and post-sessions, indicating a reduction in overall mood disturbance. (B) Changes in TMD scores for the RUN and CTL conditions. The decrease in TMD score was significantly greater in the RUN condition than in the CTL condition. (C) TA scores before and after the RUN and CTL conditions. Moderate-intensity running significantly reduced TA scores between pre- and post-sessions. (D) Changes in TA scores for the RUN and CTL conditions. The decrease in TA score was significantly greater in the RUN condition than in the CTL condition. (E) VA scores before and after the RUN and CTL conditions. (F) Changes in VA scores for the RUN and CTL conditions. The increase in VA score was significantly greater in the RUN condition than in the CTL condition. **Note**: \**p* < 0.05, \*\**p* < 0.01. In panels A, C, and E, bars represent means and error bars indicate 95% confidence intervals. In panels B, D, and F, box-and-whisker plots are drawn in the Tukey manner, overlaid error bars indicate mean ± 2 SD, dashed horizontal lines indicate zero, and line plots represent individual data. RUN = moderate-intensity running condition; CTL = seated resting control condition; TMD = total mood disturbance; TA = tension–anxiety; VA = vigor–activity.

Significant condition × time interactions were also observed for the POMS subscale scores of TA and VA (see Supplemental Table 2, Supplemental Digital Content 1, which presents repeated-measures ANOVA results for POMS subscale scores). For TA, Bonferroni-corrected post hoc comparisons showed that the score significantly decreased between the pre- and post-sessions in the RUN condition, whereas no significant pre–post change was observed in the CTL condition (Figure 3C). TA decreased significantly more in the RUN condition than in the CTL condition (*t* (26) = −2.49, *p* = .02, *d* = −0.48) (Figure 3D). For VA, a significant condition × time interaction was also observed, although Bonferroni-corrected post hoc comparisons did not reach statistical significance (Figure 3E). Nevertheless, VA increased significantly more in the RUN condition than in the CTL condition (*t* (26) = 2.49, *p* = .02, *d* = 0.48) (Figure 3F). These findings indicate that moderate-intensity running reduced TMD and TA, while increasing VA, relative to seated rest.

### Correlation analyses of running-related changes in interoception and mood states

We examined whether individual differences in running-related changes in interoception were associated with changes in mood states (see Supplemental Figure 1, Supplemental Digital Content 1, which shows exploratory correlations between running-related changes in interoceptive measures and mood scores). Running-related changes were quantified as contrast scores, calculated as the pre–post change in the RUN condition minus the pre–post change in the CTL condition (ΔRUN − ΔCTL). Greater running-related increases in BPQ scores were significantly associated with greater reductions in TMD scores (*r* = −.61, *p* < .001; Figure 4A), TA scores (*r* = −.45, *p* = .018; Figure 4B), and DD scores (*r* = −.49, *p* = .009; Figure 4C). In addition, greater running-related increases in HTT *d′* were significantly associated with greater increases in VA scores (*r* = .56, *p* = .003; Figure 4D). These findings suggest that greater running-related increases in present-moment interoceptive sensibility were associated with greater reductions in negative mood states, whereas greater increases in heartbeat-signal discriminability were associated with greater increases in positive mood.

**Figure 4.**
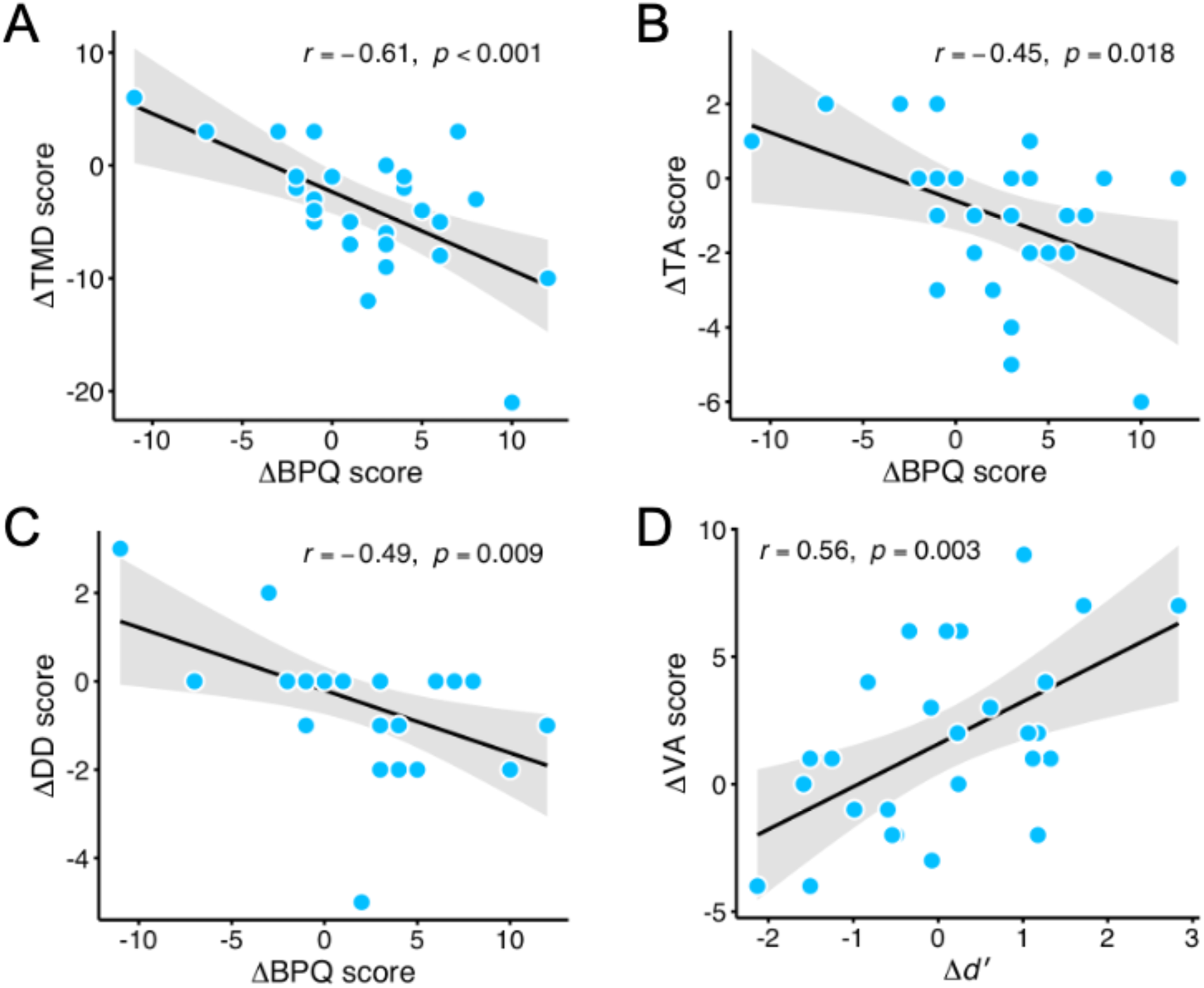
Correlations between running-related changes in interoception and mood states. (A) Association between running-related changes in BPQ score (ΔRUN − ΔCTL) and TMD score (ΔRUN − ΔCTL). (B) Association between running-related changes in BPQ score (ΔRUN − ΔCTL) and TA score (ΔRUN − ΔCTL). (C) Association between running-related changes in BPQ score (ΔRUN − ΔCTL) and DD score (ΔRUN − ΔCTL). (D) Association between running-related changes in HTT *d′* (ΔRUN − ΔCTL) and VA score (ΔRUN − ΔCTL). **Note**: BPQ = Body Perception Questionnaire; TMD = total mood disturbance; TA = tension–anxiety; DD = depression–dejection; VA = vigor–activity; HTT = heartbeat tapping task. Running-related changes were calculated as contrast scores (ΔRUN − ΔCTL), where Δ represents post–pre change. In all panels, the black line represents the linear regression line, and the gray band represents the 95% confidence interval.

### Mediation analysis of interoceptive sensibility and mood

We examined whether changes in present-moment interoceptive sensibility statistically accounted for the effect of moderate-intensity running on mood improvement (Figure5A). Because changes in the mediator and outcome were assessed over the same pre–post interval, these mediation analyses were interpreted as preliminary and as testing whether the data were statistically consistent with an indirect pathway. Mediation analyses showed that changes in BPQ score partially mediated the effect of running on reductions in TMD score (ACME = −1.36, 95% CI [−3.04, −0.01], *p* = .04; ADE = −2.33, 95% CI [−4.11, −0.54], *p* = .01; total effect = −3.69, 95% CI [−5.82, −1.56], *p* = .002; proportion mediated = 0.36, 95% CI [0.03, 0.78], *p* = .04) (Figure 5B). These findings suggest that increased present-moment interoceptive sensibility may partly account for the reduction in TMD after running.

**Figure 5.**
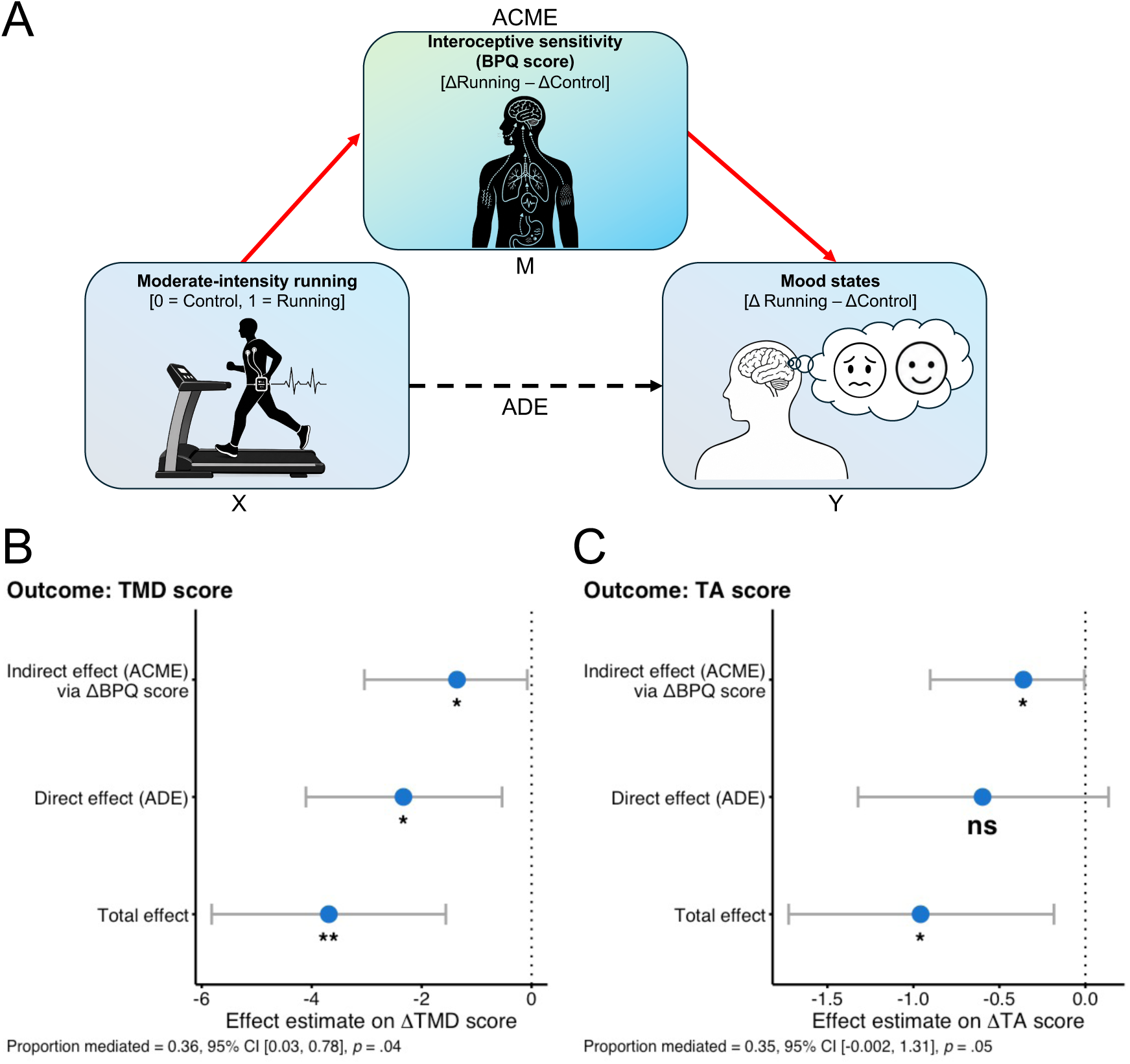
Mediation analysis of interoceptive sensibility and mood changes. (A) Mediation model testing whether changes in present-moment interoceptive sensibility, indexed by Body Perception Questionnaire (BPQ) score, statistically accounted for the effect of moderate-intensity running on changes in mood states. For each condition, change scores were calculated as post-session minus pre-session scores, and the exposure variable was condition (0 = CTL, 1 = RUN). (B) Results of mediation analysis for total mood disturbance (TMD), the primary mood outcome. Changes in BPQ score showed a significant average causal mediation effect (ACME), while the average direct effect (ADE) also remained significant, suggesting that the data were statistically consistent with partial mediation. (C) Results of exploratory mediation analysis for tension–anxiety (TA). Changes in BPQ score showed a significant ACME, whereas the ADE was not significant and the proportion mediated did not reach conventional statistical significance; therefore, this indirect association should be interpreted as modest and exploratory. Note: BPQ = Body Perception Questionnaire; TMD = total mood disturbance; TA = tension–anxiety; RUN = moderate-intensity running condition; CTL = seated resting control condition; ACME = average causal mediation effect; ADE = average direct effect. In panels B and C, points represent estimated effects, and error bars indicate 95% confidence intervals. \**p* < .05, \*\**p* < .01; ns, not significant.

We also conducted an exploratory mediation analysis for TA, given its significant association with running-related changes in BPQ score. Changes in BPQ score showed a significant indirect association between running and reductions in TA score (ACME = −0.36, 95% CI [−0.90, −0.01], *p* = .046). However, the ADE was not significant (ADE = −0.60, 95% CI [−1.32, 0.13], *p* = .12), and the proportion mediated did not reach conventional statistical significance (proportion mediated = 0.35, 95% CI [−0.002, 1.31], *p* = .05) (Figure 5C). Thus, the mediation result for TA should be interpreted as modest and exploratory.

## Discussion

The present study examined whether acute moderate-intensity running changes interoception and mood states, and whether running-related changes in interoception are associated with mood improvement. The main findings were as follows. First, moderate-intensity treadmill running increased self-reported present-moment interoceptive sensibility, as indexed by BPQ scores, but did not significantly change task-based measures of cardiac interoceptive accuracy, including HCT score and HTT-derived md, mSI, and *d′*. Second, running reduced TMD and TA and increased VA compared with seated rest. Third, greater running-related increases in BPQ scores were associated with greater reductions in TMD, TA, and DD. Greater running-related increases in HTT *d′* were also associated with greater increases in VA. Finally, mediation analyses suggested that changes in BPQ score partly accounted for the effect of running on reduced TMD, with weaker and exploratory evidence for TA. These findings suggest that subjective awareness of bodily sensations may be one process related to mood improvement after acute running.

The present study found that moderate-intensity running improved mood states, as reflected by reduced TMD and TA and increased VA. This pattern is consistent with previous evidence showing that acute aerobic exercise can reduce anxiety and improve affective states (1,4). It is also consistent with running-specific findings showing that even a brief bout of moderate-intensity running can increase pleasant mood and arousal (28,29). In the present study, the RUN condition was performed at 50% of estimated V̇O_₂peak_ and lasted 15 min, indicating that a relatively short bout of moderate-intensity running was sufficient to improve both negative and positive mood states. These findings support the practical value of moderate-intensity running as a simple exercise mode for improving transient mood states in healthy young adults.

A key finding of the present study is that running-related increases in BPQ scores were associated with improvements in mood states and statistically accounted for part of the effect of moderate-intensity running on TMD. A similar, although more modest and exploratory, pattern was observed for TA. These findings are consistent with the possibility of an exercise–interoception–mood pathway, in which exercise-induced changes in subjective interoceptive sensibility contribute to mood improvement. Physical activity has been proposed to increase visceral-afferent input and the salience of bodily signals entering the interoceptive system, thereby influencing how bodily changes are processed and integrated with cognitive, motivational, and affective responses (11,16,20,55). Thus, acute running may have produced salient bodily changes, including increased HR and respiration, and individuals who became more aware of these changes may have been better able to integrate them into subjective emotional experiences. This interpretation is also consistent with interoceptive exposure accounts, which suggest that salient physiological arousal induced by physical activity may increase familiarity with bodily arousal cues and reduce catastrophic or negative interpretations of these sensations (56). Accordingly, subjective interoceptive sensibility may be one proximal process through which moderate-intensity running contributes to improvements in mood states (Figure 6), although it likely operates alongside other physiological and psychological mechanisms, such as neurochemical changes, autonomic regulation, self-efficacy, and attentional distraction (4,9).

**Figure 6.**
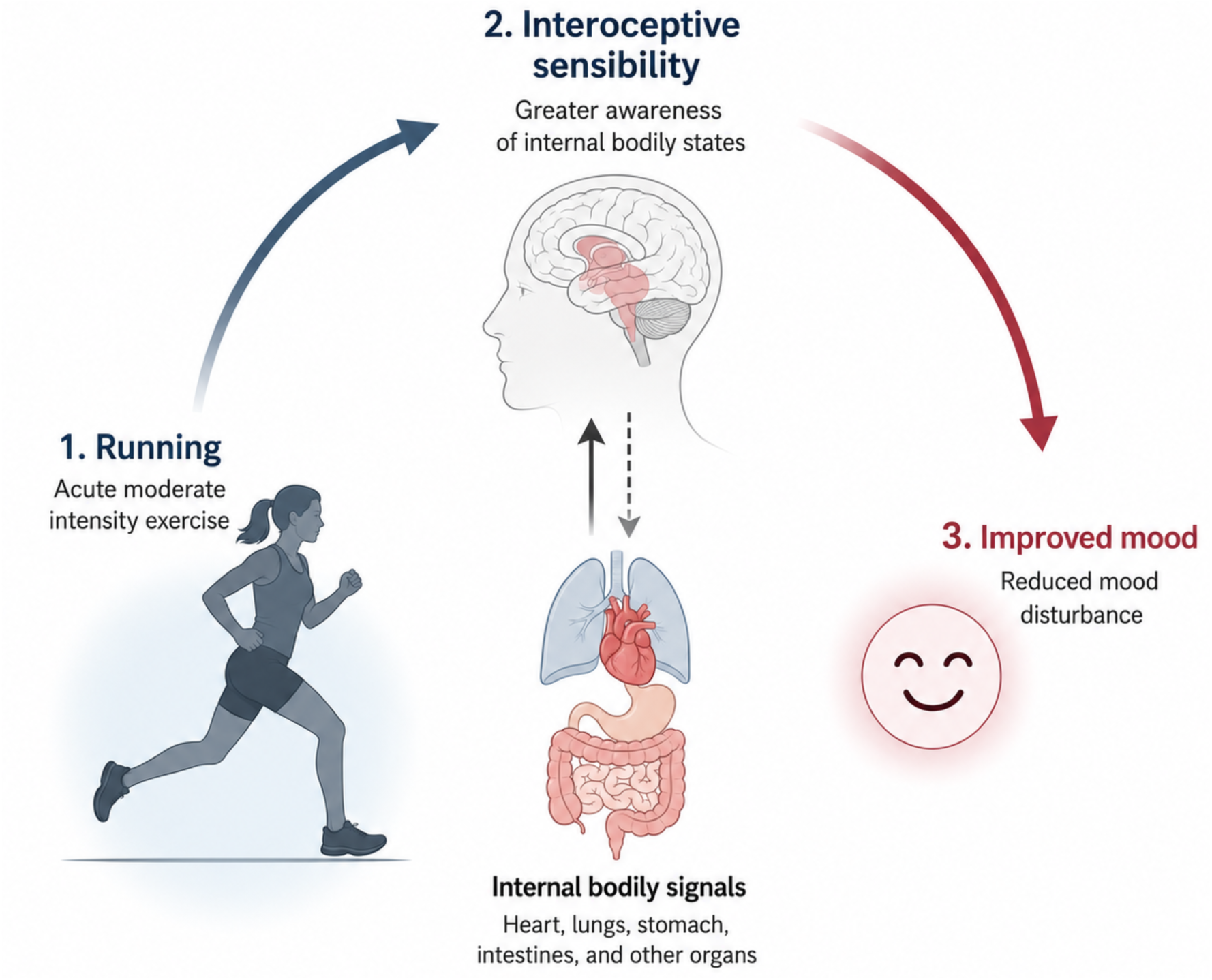
Conceptual schematic of the proposed running–interoception–mood pathway. Conceptual schematic illustrating the proposed pathway suggested by the present findings. Acute moderate-intensity running may increase present-moment interoceptive sensibility, reflected as greater awareness of internal bodily states. Internal bodily signals arising from organs such as the heart, lungs, stomach, and intestines are represented as contributing to this interoceptive sensibility. Increased interoceptive sensibility may, in turn, be related to reduced mood disturbance. This schematic is intended as a conceptual summary of the present results and does not imply that all depicted pathways were directly tested.

In contrast to our initial expectation, moderate-intensity running did not significantly change task-based indices of interoceptive accuracy. No significant condition × time interactions were observed for HCT score or for the HTT-derived indices of md, mSI, and *d′*. This suggests that running did not produce a robust group-level improvement in cardiac interoceptive accuracy in the present study, despite increased cardiac activity after running. This finding differs from previous studies showing that cardiac awareness or heartbeat perception performance can improve during or after exercise (21,22,23). However, those studies involved stronger exercise-induced cardiac activation, assessment during exercise, or higher-intensity physical activity than the present protocol. Thus, one possible explanation is that the bodily signals induced by the present moderate-intensity running protocol may not have been strong enough, or sufficiently specific to cardiac sensations, to improve task-based heartbeat perception. This interpretation is also consistent with the idea that improvement in heartbeat perception performance may require sufficiently increased salience of cardiac sensations. For example, Smith et al. (17) showed that healthy participants improved heartbeat tapping performance during an inspiratory breath-hold condition designed to amplify cardiac sensations. Breath holding may improve heartbeat perception performance relatively directly by increasing the salience of cardiac sensations. In contrast, the post-running state is likely to involve a broader set of exercise-related bodily cues, including cardiovascular, respiratory, thermal, muscular, and effort-related sensations (57,58). Such broad exercise-related bodily cues may influence affective responses but may not necessarily enhance cardiac sensations in a sufficiently specific or direct manner to improve heartbeat perception performance. Thus, differences in exercise intensity, the specificity of bodily manipulation, and assessment timing may help explain why task-based interoceptive accuracy did not improve in the present study.

Although moderate-intensity running did not produce a significant group-level increase in HTT *d′*, the exploratory association between running-related changes in HTT *d′* and VA suggests that individual differences in cardiac signal discrimination may be related to positive activation-related mood responses. Specifically, individuals who showed greater increases in *d′* also showed greater increases in vigor–activity. One possible interpretation is that individuals who became more sensitive to cardiac signals after running may have experienced exercise-induced bodily activation as a more salient and energizing state. In this sense, changes in HTT *d′* may not reflect a general improvement in interoceptive accuracy, but rather an individual difference in how cardiac bodily signals are integrated into positive mood states. This possibility is consistent with broader accounts proposing that interoceptive processing contributes to emotional experience and affective regulation (12,20). However, because this association was exploratory and the present study did not directly manipulate cardiac signal discrimination, this interpretation should be regarded as preliminary.

The present findings may also be considered in relation to predictive processing or active inference accounts of interoception and emotion, although the present study was not designed to directly test these frameworks. From this perspective, emotional experience is not determined solely by peripheral bodily changes themselves but by how the brain interprets, weighs, and updates internal bodily signals (59,60). Moderate-intensity running produces clear and salient changes in bodily states, such as increased HR and respiration. Importantly, these bodily changes occur in a context in which their causes are identifiable and predictable, namely the physiological demands of physical activity. One speculative interpretation is that exercise-induced bodily signals may provide relatively clear interoceptive evidence for updating the subjective representation of bodily states, rather than remaining ambiguous internal sensations. This framework may help explain why changes in BPQ scores, but not all indices of interoceptive accuracy, were associated with mood improvement: what may be critical is not simply detecting cardiac signals more accurately, but becoming subjectively aware of bodily changes in a context in which those signals can be meaningfully interpreted and integrated into emotional experience (12,20). However, because the present study did not directly measure bodily appraisal, neural prediction errors, precision weighting, or brain activity, this interpretation should be regarded as a theoretical account rather than direct evidence for predictive coding mechanisms (60,61).

In conclusion, a single bout of moderate-intensity treadmill running improved mood states and increased present-moment interoceptive sensibility in healthy young adults. Specifically, running reduced TMD and TA, increased VA, and increased BPQ scores, but did not significantly alter task-based indices of cardiac interoceptive accuracy. Running-related increases in BPQ scores were associated with reductions in negative mood states and were statistically consistent with an indirect pathway linking running to reduced TMD. These findings suggest that enhanced subjective interoceptive sensibility may be one proximal process related to the mood benefits of acute running. Together, the present findings support the possibility of an exercise–interoception–mood pathway, in which exercise-induced bodily changes are linked to subjective mood states through increased awareness of bodily sensations.

## Acknowledgements

The authors thank all participants for their involvement in this study.

## Data availability

The de-identified datasets supporting the findings of this study and the analysis code used to reproduce the main statistical analyses and figures are available from the corresponding author upon reasonable request. Data and code will be shared for research purposes, where appropriate, in accordance with institutional and ethical requirements.

## Supplemental Digital Content

Supplemental materials are available at https://osf.io/h4yns/files/dfahv.

